# Correspondence on “Fortification of FeS Clusters Reshapes Anaerobic CO Dehydrogenase into an Air-Viable Enzyme Through Multilayered Sealing of O_2_ Tunnels”

**DOI:** 10.64898/2026.02.25.707409

**Authors:** Laura V. Opdam, Peter Gebhardt, Christophe Léger, Holger Dobbek, Vincent Fourmond

## Abstract

In their recent communication in Angewandte Chemie (10.1002/anie.202508565), Suk Min Kim and coworkers have described the effect of modifying the gas channels of the CO dehydrogenase II from *Carboxydothermus hydrogenoformans*, an enzyme that oxidizes reversibly CO into CO_2_. Their goal was to use mutagenesis to slow down the arrival of O_2_ at the active site. They reported a large increase in the resistance against oxygen, one of the major barriers to the application of this extremely fast and efficient enzyme in biotechnological devices, with an increase in the IC50 of more than two orders of magnitudes for some variants, with only a minor impact on the affinity of the enzyme for CO. We have produced the same variants, and characterized them in depth using Protein Film Electrochemistry. We used an approach that has proven very useful to learn and understand about the reactivity of CO dehydrogenases (and other redox enzymes like hydrogenases) with O_2_. We found that, contrary to the claims by Kim and coworkers, the A559W and the A559W/V610H mutants are not more resistant than the WT against oxygen.

## Introduction

NiFe-containing CO dehydrogenases (CODH) are the enzymes that catalyze the reversible reduction of CO_2_ to CO, at a unique NiFe_4_S_4_ active site^[1–3]^. They are known for their very high catalytic rates^[4]^ and also for their high energetic efficiency^[5]^, which makes them particularly interesting for incorporation into biochemical devices like bioelectrolyzers or biosensors. These possibilities however are hindered by the high sensitivity of these enzymes to molecular oxygen, greatly complicating the construction and operation of CODH-based devices. The reactivity of CODHs with O_2_ is not well understood. We have shown that O_2_ targets the active site^[6]^, and that the behaviour of the enzymes varies significantly from one CODH to another: for instance, the CODH from *Nitratodesulfovibrio vulgaris* (formerly *Desulfovibrio vulgaris*) inactivates at very low concentrations of O_2_, but is reactivated upon reduction^[6]^, as a result of a significant O_2_-induced conformational change of the active site^[7]^, while *Carboxydothermus hydrogenoformans* CODH IV resists to much higher concentrations of O_2_, but react irreversibly with O_2_^[8]^. We have also been able to elucidate the first steps of the O_2_-induced degradation of the C cluster of *C. hydrogenoformans* CODH II (CODH-II_*Ch*_)^[9]^. Wang and coworkers have also used electrochemistry to characterize the reactivity of CODH-II_*Ch*_ with O_2_; they were able to show that high potentials and CN^-^ protect against O_2_ damage^[10]^.

The active site of CODHs is buried within the protein matrix, with gas channels that connect it to the outside to permit the intramolecular diffusion of CO and CO_2_^[11,12]^; presumably, these channels also guide the diffusion of O_2_ towards the active site, hence the intuitive approach of blocking the gas channel to increase the resistance to O_2_. Some of us have previously employed this strategy with NiFe hydrogenases, using a systematic mutagenesis approach to study the influence of the amino acid introduced in the gas channel on the diffusion of H_2_, O_2_ and CO^[13]^. This led us to learn how the modifications of the gas channel affect diffusion rates, and to conclude that CO and O_2_ diffuse within the protein matrix of hydrogenase at the same rate^[13]^.

In a recent Angewandte Chemie Research Article^[14]^ based on a previous mutagenesis work^[15]^, Kim and coworkers report on a series of single and double replacements of residues in one of the branches of the gas channel of CODH-II_*Ch*_ that increase the O_2_ half maximum inhibitory concentration (IC50) of the enzyme by more than two orders of magnitude, when measured using solutions assays. The strategy was to introduce bulky residues to replace alanine 559, an amino acid in the vicinity of the active site, and valine 610, which is on the surface (figure 1). Among the mutants tested in this work, the A559W/V610H double mutant showed the highest resistance against oxygen (a factor of 300 increase in IC50), with almost no impact on the *K*_*M*_ for CO.

**Figure 1.**
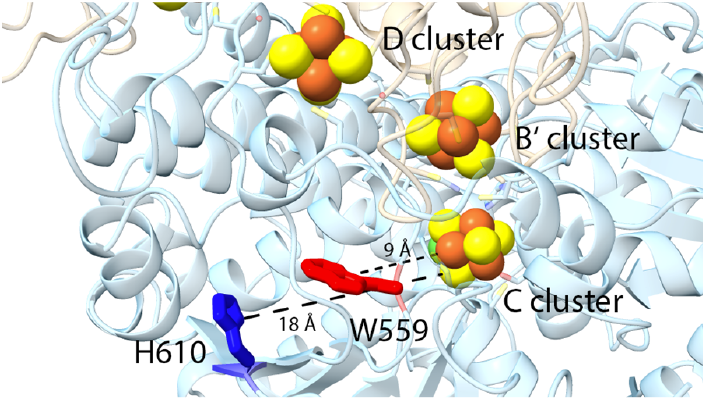
Position of exchanged residues in the A559W/V610 variant. The two residues W559 and H610 are highlighted in red and blue respectively. The shortest distances between the residues and the nearest C-cluster atoms are displayed.

To better understand the origin of this alleged high resistance, we constructed, expressed and purified the single A559W and the double A559W/V610H mutants of CODH-II_*Ch*_, and characterized them using protein film electrochemistry^[16,17]^. In this technique, the enzyme is immobilized on an electrode in a configuration of direct electron transfer, and the catalytic activity is measured as a current and monitored with a high time resolution (0.1 s in our experiments). We have previously developed a strategy to submit the immobilized enzyme to transient exposures to O_2_^[18]^ (of about 30 to 50 s), and have used it to characterize in detail the reactivity with oxygen of a number of CODHs: *Nitratodesulfovibrio vulgaris (Nv)* CODH, CODH-II_*Ch*_^*[6]*^*C. hydrogenoformans* CODH IV^[8]^ and *Thermococcus sp. AM4* CODH 1 & 2^[19]^. This technique has proven more sensitive and more insightful than solution assays, because i) the rotation leads to fast homogenization of the injected oxygen (about 0.1 s) and ii) the technique allows for a constant monitoring over time of the remaining activity.

In contrast to the original article, we found that the A559W and the A559W/V610H mutations have no impact on the sensitivity of the enzyme to oxygen.

## Results

We expressed and purified WT CODH-II_*Ch*_ and the two mutants according to procedures established in the group from Berlin^[20]^. As a quality control and a proof that the mutations were indeed introduced, we crystallized the mutants and determined their structures, showing that indeed the mutations were properly introduced (Supplementary Figures 1 and 2). The crystal structures are described in supplementary information section S1-2, along with the details about the mutagenesis. The structures were deposited in the Protein Database with the accession codes 9TOX and 9TPO.

Over the past 10 years, we have adapted to the study of CODH electrochemical methods that were initially developed to examine the reactions of hydrogenases with gaseous substrate and inhibitors^[21–23]^: the catalytic current (and hence the catalytic activity) is followed as a function of time after the injection of an aliquot of CO or O_2_-saturated solution in the electrochemical cell; due to the exchange with the atmosphere above the electrochemical cell, their concentrations decrease exponentially over time, and the time-dependent current reports on how the activity depends on the concentration of substrate and inhibitor (Figure 2).

**Figure 2.**
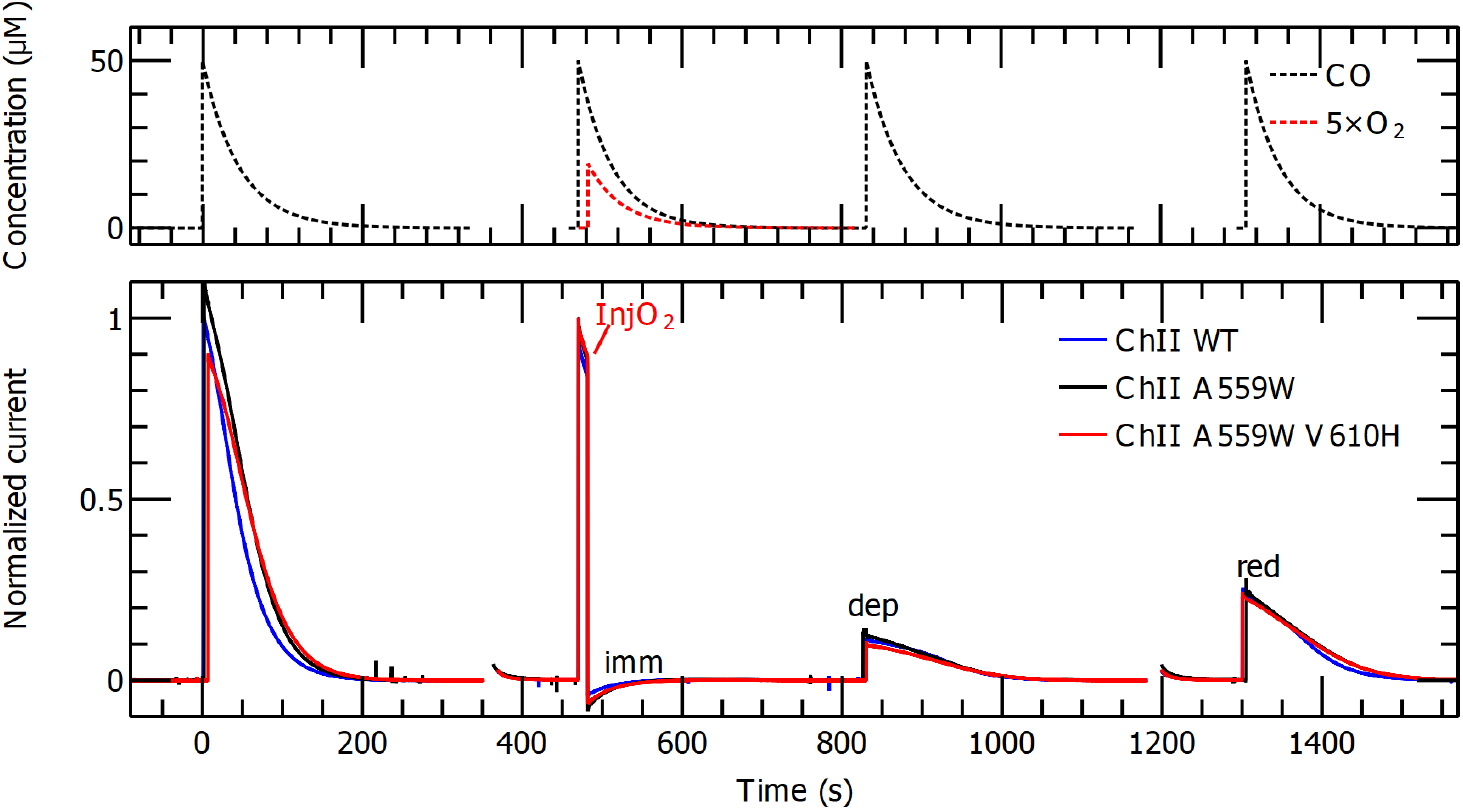
One of the experiments used to characterize the reactivity of the WT and the two mutants with O_2_. Top: concentrations of CO and O_2_ over time. Bottom: response in current of CODH-II_*Ch*_ WT and the two variants, normalized to account for different electroactive coverages.

Using this approach, we have determined in this work the Michaelis constants of the WT CODH-II_*Ch*_ and the two variants by exposing films of the enzymes to injections of CO (Supplementary Figure 3). The exponential decrease in CO concentration as a function of time makes it possible to explore a large range of concentrations in a single experiment; appropriate modeling^[24]^ yields the value of the Michaelis constant (table 1), which increases moderately in the order WT < A599W < A559W/V610H. These results are compatible with those of Kim and coworkers, who found respectively 20±2 µM, 34±2 µM and 90±20 µM^[14]^: although the absolute values are smaller in electrochemistry, the impacts of the mutations in relative terms are very similar between the work of Kim and coworkers and ours.

**Table 1.** The concentration of oxygen needed to inactivate 50% of the enzyme (IC50) directly after oxygen injection K ^imm^, after oxygen departed from the electrochemical cell K^dep^, and after a reductive poise K ^red^. The procedure used to determine the value and the error of the IC50 values is presented in supplementary section S4. The value of the K_M_, which is an average of at least 3 measurements taken on at least 2 separate days.

| Variant | $K_i^{\text{imm}}$ ($\mu\text{M O}_2$) | $K_i^{\text{dep}}$ ($\mu\text{M O}_2$) | $K_i^{\text{red}}$ ($\mu\text{M O}_2$) | $K_M$ ($\mu\text{M CO}$) |
| --- | --- | --- | --- | --- |
| CODH-II<sub>Ch</sub> WT | 0.03±0.02 | 0.05±0.02 | 0.3 ± 0.1 | 4.5±0.8 |
| CODH-II<sub>Ch</sub> A559W | 0.03±0.02 | 0.07±0.02 | 0.3 ± 0.1 | 7.7±0.9 |
| CODH-II<sub>Ch</sub><br>A559W/V610H | 0.03±0.02 | 0.08±0.02 | 0.24 ± 0.06 | 11±2 |
| CODH<sub>Nv</sub> | 0.03±0.02 | 3±1 | >30 | 3±1 |

To assess the reactivity of the CODHs with O_2_, we used a previously described protocol combining several injections of CO to measure the initial and remaining activities and one injection of O_2_^[6]^. Figure 2 shows a typical experiment (for the WT and the two variants), with four injections of CO (at t=0, 470, 840 and 1300 s) and one injection of O_2_ at t = 480 s (the full experiment includes two other CO injections, which help to take into consideration film loss, see more details in ref. ^[6]^). The top panel represents the concentrations of CO and O_2_, and the bottom panel shows the current response of the three enzymes. Each injection of CO results in the appearance of a CO oxidation current that decreases over time because of the decrease in CO concentration. Just after the second injection of CO, the O_2_ injection results in a fast decrease in current indicating a fast inactivation step (here leading to a complete loss of catalytic current, the small negative current results from direct O_2_ reduction at the electrode). The presence of a CO oxidation current after the third injection shows that the enzyme has reactivated partially after the departure of O_2_, and a further increase in current at the fourth injection (following the reductive poise) shows that reduction has further reactivated the enzyme. From the data shown in figure 2, it is very difficult to see any difference between the WT and the two mutants: the loss in activity just after the O_2_ injection and the activities recovered after the departure of O_2_ and after the reductive poise are the same for all variants.

To further compare the WT and the mutants, we repeated this experiment, but varying the quantity of injected O_2_ (Figure 3). From each experiment, it is possible to determine three remaining activities: the one immediately after the O_2_ injection (“imm”, blue symbols), after O_2_ departure (“dep”, black symbols) and after the reductive poise (“red”, red symbols). These values are plotted in figure 3A as a function of the injected O_2_ concentration for the WT and the two mutants. The data show very little difference between the three enzymes. As a positive control, we have included in figure 3B the data that were acquired with *Nitratodesulfovibrio vulgaris (Nv)* CODH using the same method: they show little difference in the “imm” response, but very important differences for “dep” and “red”: as was reported earlier, CODH_*Nv*_ is able to reactivate significantly upon reduction after it has been inhibited by O_2 ^[6]^_.

**Figure 3.**
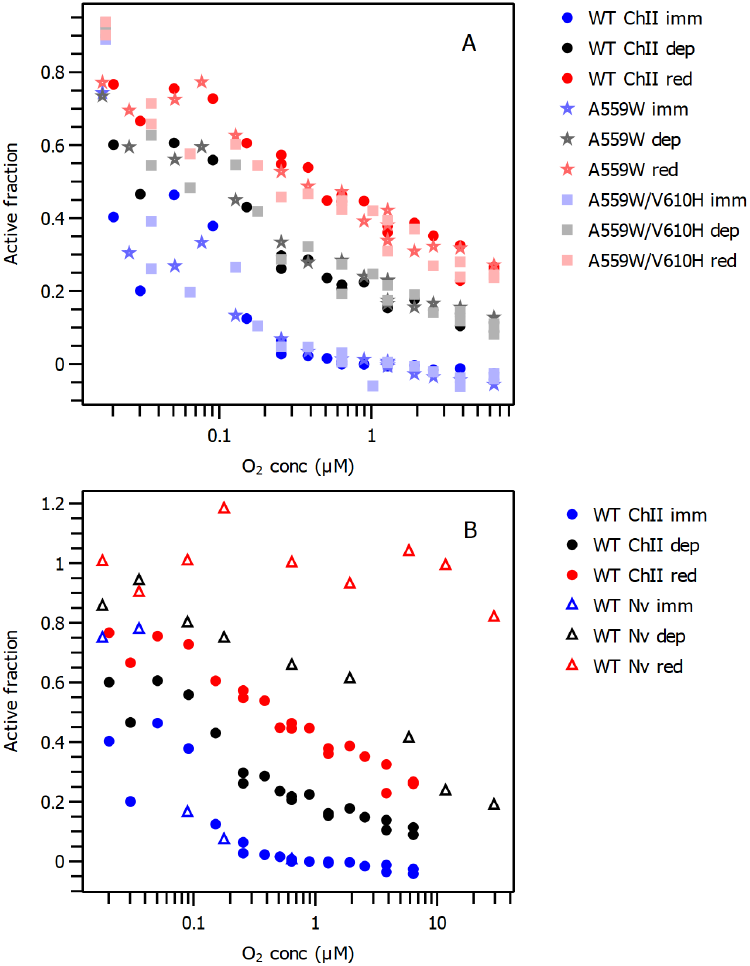
The fraction of remaining active enzyme measured just after an injection of oxygen (imm), after oxygen departure from the electrochemical cell (dep) and after a reductive poise (red), plotted as a function of the initial concentration of oxygen. A: data for *Ch* CODH II WT (circles), A559W (stars) and A559W/V610H (squares). B: data for *Ch* CODH II WT (circles, same data as panel A), and *Nv* CODH (open triangles, data replotted from ref ^[6]^).

We used the method described in the SI to determine the corresponding IC50 values for the different curves, i.e. the injected O_2_ concentration that inactivates 50% of the enzyme; the results are given in table 1. In line with the almost complete superimposition of the curves in figure 2, the values of the various IC50 for the WT and the two mutants are not significatively different. On the other hand, the IC50 values determined for CODH_*Nv*_ are much greater than for WT CODH-II_*Ch*_.

## Discussion/conclusion

We have used state-of-the-art protein film electrochemistry methods to characterize the reactivity of CODH-II_*Ch*_ and its two variants A559W and A559W/V610H with CO and with oxygen. We found only small differences in the reactivity of the enzymes with CO, with up to a factor of 3 increase in the values of *K*_*M*_. We found no significant difference in the IC50 for the inhibition of the CODHs by oxygen. While the increases in the values of *K*_*M*_ that we determined are consistent with what Kim and coworkers measured^[14]^, the data of the reactivity with oxygen here clearly contradict the results reported by Kim and coworkers^[14]^, who found that the values of the IC50 were increased 50-fold for the A559W mutant, and around 300-fold for the A559W/V610H mutation. The origin of the discrepancy is unclear to us. The electrochemical approach we have used here has been validated with a number of CODHs before, and the insights obtained using electrochemistry were confirmed by other methods: for instance, the observation by electrochemistry that *N. vulgaris* CODH reactivates after a reductive poise was explained by the crystal structure of an alternative form of the CODH active site after exposure to O_2_, which reverts to the canonical structure after reduction^[7]^.

The lack of significant improvement of the oxygen resistance of CODH-II_Ch_ in the two mutants is not surprising. Analysis of crystal structures (including with xenon binding) show the existence of a series of gas channels that converge into a single branch near the active site^[11]^, which is similar to what has been observed in other enzymes like NiFe and FeFe hydrogenases^[25–27]^. As a consequence, it is expected that only mutations that are close enough to the active site to target the last common channel would impair significantly the access of gaseous molecules towards the active site. Here, A559 is 8.7 Å away from the Ni of the active site, past the first fork in the channel, and V610 is 18.6 Å away, at the surface of the protein. These observations suggest that the mutations should only have a small impact on the diffusion of CO (as indeed observed both by us and by Kim and coworkers), and, similarly, a small impact on the diffusion of O_2_, which is clearly demonstrated here but incompatible with the reported results of Kim and coworkers.

## Supporting Information

The supporting information contains the molecular biology/biochemistry procedures, the crystal structures of the mutants, details about the experimental procedures, in particular the determination of the KM and the IC50.

The raw electrochemical data were deposited on the Zenodo repository 10.5281/zenodo.18468844.

## Acknowledgements

The authors acknowledge support from CNRS, Agence Nationale de la Recherche (grants ANR-17-CE11-0027, ANR-21-CE50-0041, ANR-23-SODR-0004, ANR-23-CE44-0046), and Region PACA. We further acknowledge funding by the Deutsche Forschungsgemeinschaft (DFG) through grant DO 785/10-1 and access to the BESSY II storage ring (Berlin) through the Joint Berlin MX-Laboratory sponsored by Helmholtz-Zentrum Berlin für Materialien und Energie, Freie Universität Berlin, Humboldt-Universität zu Berlin, Max-Delbrück-Centrum and the Leibniz-Institut für Molekulare Pharmakologie. This project has received funding from the European Union’s Horizon Europe research and innovation programme under grant agreement number 101115403 (ECOMO). This work has benefited from French State aid managed by the Agence Nationale de la Recherche under France 2030 plan, bearing the reference code ANR-22-PESP-0010: Projet ciblé “POWERCO2” within the PEPR project SPLEEN. The project leading to this publication has received funding from Excellence Initiative of Aix-Marseille University—A*Midex, a French “Investissements d’Avenir” program. LVO, CL and VF are members of the French Bioinorganic Chemistry group (http://frenchbic.cnrs.fr).

