## Supplementary information for "Correspondence on “Fortification of FeS Clusters Reshapes Anaerobic CO Dehydrogenase into an Air-Viable Enzyme Through Multilayered Sealing of O_2_ Tunnels”"

Laura V. Opdam[a], Peter Gebhardt[b], Christophe Léger[a], Holger Dobbek[b]* and Vincent Fourmond[a]*

[a] LVO, CL and VF
Laboratoire de Bioénergétique et Ingénierie des Protéines. CNRS, Aix Marseille Université, UMR, 7281 Marseille, France


[b] PG and HD
Institute of Biology, Humboldt-Universität zu Berlin, Philippstraße 13, 10115, Berlin, Germany

**S1 Plasmid preparation and protein purification** 1
**S2 Crystallography** 2
**S3 Electrochemistry** 4
**References** 5

### S1 Plasmid preparation and protein purification

The plasmid pKS2 containing the *cooS-II*-gene encoding CODH-II$_{Ch}$ was amplified using mismatch primers to generate site-directed nucleotide exchanges. For CODH-II$_{Ch}$ A559W the following primers were used: 5'-CTGCCGAATGGATGCATGAAAAG-3' and 5'-CAGAAGCTACAACCGGAAGTCG-3'. For the double variant CODH-II$_{Ch}$ A559W-V610H the following primers were used on template DNA including the prior exchange of A559W: 5'-CTTTATTCACGAACTTGATCCCG-3' and 5'-TAGCCTCCGGTTATATCTTTG-3'. For PCR the Q5-Mastermix (New England Biolabs) was used. The PCR product was treated with *Dpn I* (Thermo Fisher Scientific), phosphorylated with T4 polynucleotide kinase (Thermo Fisher Scientific) and ligated using T4 DNA ligase (Thermo Fisher Scientific), mimicking the KLD Enzyme Mix Reaction Protocol (New England Biolabs #M0554). After transformation into *E. coli* XL1 blue (DE3) cells, the successful mutagenesis was confirmed by DNA sequencing, using DNA prepared from visible colonies (NucleoSpin Plasmid Kit, Macherey-Nagel).

For protein expression, we used *E. coli* BL21 (DE3) cells containing the pRKISC plasmid[1]. The cells were grown in modified terrific broth medium at 30° C under constant stirring and flushing with air. The medium was supplemented with 1% (w/v) glucose, 2 mM L-cysteine, 0.6 mM $FeSO_4$, 0.04 mM $NiCl_2$ and 0.45 mM NaS. Once the culture had reached an optical density of 0.7 at 600 nm, flushing was switched to nitrogen and expression was induced by adding 0.2 mM IPTG. After induction 50 mM $KNO_3$, 1 mM $FeSO_4$, 0.95 mM $NiCl_2$ and 0.75 mM $Na_2S$ were added to the medium. Expression continued at 30° C and cells were harvested after 20 - 22 h. The cell pellets were frozen in liquid nitrogen and stored at -80° C until purification.

For purification, all steps were carried out inside an anoxic glove box (Model B, COY Laboratory Products). The atmosphere inside the glove box consisted of 95% $N_2$ and 5% $H_2$ at 20 °C. We followed the steps as previously described[1]. The details of this process can be seen below.

The cell pellets were resuspended in lysis buffer (50 mM Tris-HCl pH 8.0, 20 mM NaCl, 20 mM imidazole, 10% (w/v) saccharose), supplemented with DNase and lysozyme and stirred for 20 min. Afterwards, we lysed the cells by sonification in two steps for 5 min each on ice. After adding 0.9% (v/v) deoxycholic acid, the lysate was stirred for 1 h and then centrifuged at 21,000 g for 20 min. The supernatant was loaded onto a $Ni^{2+}$-immobilized metal affinity chromatography column (Ni Sepharose, GE Healthcare), which was equilibrated with wash buffer (50 mM Tris-HCl pH 8.0, 20 mM NaCl, 20 mM imidazole). After washing the loaded column with a wash buffer, the protein was eluted using an elution buffer containing 50 mM Tris-HCl pH 8.0, 20 mM NaCl and 250 mM imidazole. The protein-containing buffer was exchanged to G25 buffer (20 mM Tris-HCl pH 8.0) using a PD-10 desalting column (GE Healthcare). The protein was concentrated to 250 – 300 µL using a centrifugal filter unit with a 30 kDa weight cut-off (Vivaspin 500, Vivascience), transferred to a cryo vial and stored in liquid nitrogen.

Protein concentration was determined according to Bradford[2] using the Quick Start Bradford Protein Assay (Bio-Rad).

The specific CO oxidation activities of CODH-$II_{Ch}$ wildtype protein and the two produced variants were determined as previously described[3].

The protein was first diluted with a buffer (20 mM Tris-HCl pH 8.0, 3 mM Na-dithionite, 2 mM dithiothreitol). Then the protein was injected into a quartz cuvette containing the activity buffer (50 mM HEPES-NaOH pH 8.0, 20 mM oxidized methyl viologen, 2 mM dithiothreitol) with CO headspace. The reduction of oxidized methyl viologen was monitored at 578 nm ($\varepsilon_{578}$ = 9.7 $mM^{-1}$ $cm^{-1}$ of reduced methyl viologen) and 22º C. Each measurement was repeated three times and the average value, together with its standard deviation is reported here. The calculated specific CO oxidation activity of wildtype CODH-$II_{Ch}$ was 1748.38 ±162.54 U $mg^{-1}$, where one unit (U) is defined as one µmol of CO oxidation per minute (this is the definition that has been used for the last 25 years, though Kim and coworkers report activities as µmol of ethylviologen reduced per minute, so that the activities they report must be divided by two for comparing with the literature). For the variant A559W it was 1547.72 ±86.63 U $mg^{-1}$ and for the variant A669W-V610H it was 1060.17 ±156.33 U $mg^{-1}$.

### S2 Crystallography

We crystallized the CODH-$II_{Ch}$ variants inside an anoxic glove box (Model B, COY Laboratory Products). Crystals were grown using hanging drop vapor diffusion over a reservoir solution containing 15 – 20% polyethylene glycol 2000 monomethyl ether (PEG 2000 MME), 100 mM sodium 4-(2-hydroxyethyl)-1-piperazineethanesulfonic acid (HEPES) and 2 mM Na dithionite at pH 7.5. The protein (16 mg/mL concentration) was diluted 1:1 with reservoir solution for a total drop volume of 4 µL. The crystals were collected for measurement in 25% (w/v) PEG 2000 MME and 100 mM HEPES and were frozen in liquid nitrogen.

Diffraction data of the protein crystals were collected at the beamline 14.1 at Helmholtz-Zentrum Berlin at the BESSY-II storage ring at a wavelength of 0.91841 Å[4]. Diffraction data were integrated and scaled with XDSAPP[5,6]. The initial phases were obtained by Patterson search methods using PhaserMR[7] with 3B51.pdb[1] as a homologous search model. After model building using coot[8] further refinements were carried out using refmac5[9] as part of the *CCP4i2* software suite[10,11]. The statistics of data collection and refinement for both CODH-$II_{Ch}$ variants are listed in Table S1.

WILEY-VCH

**Table S1.** Statistics of data collection and structure refinement of CODH-II$_{Ch}$ A559W and A559W-V610H.

| **Name** | **A559W** | **A559W V610H** |
|---|---|---|
| PDB ID | 9TOX | 9TP0 |
| **Data Collection** | | |
| Wavelength (Å) | 0.9184 | 0.9184 |
| Space Group | C2 (5) | C2 (5) |
| Cell Constants | | |
| a, b, c (Å) | 111.31, 75.33, 71.59 | 112.00, 74.98, 70.82 |
| α, β, γ (°) | 90.0, 111.56, 90.0 | 90.0, 111.15, 90.0 |
| Resolution (Å) | 40.77-1.49 (1.58-1.49)* | 40.71-1.41 (1.50-1.41)* |
| No. Reflections total/ unique reflections | 599,340/ 175,782 (28,446) | 703,541/ 104,990 (16,923) |
| Multiplicity | 3.41 | 6.7 |
| $R_{meas}$ (%) | 21.1 (135.2) | 9.9 (164.4) |
| (I)/(σI) | 9.18 (1.06) | 12.70 (1.11) |
| $CC_{1/2}$ (%) | 99.8 (32.2) | 99.9 (42.1) |
| ISa | 34.79 | 31.43 |
| Completeness (%) | 99.4 (99.8) | 99.8 (99.9) |
| **Refinement** | | |
| Nr. of reflections used in refinement | 88,496 | 104,990 |
| R/Rfree-factor (%) | 17.49/21.26 | 15.47/18.40 |
| RMSD from ideal geometry | | |
| Bonds (Å) | 0.013 | 0.016 |
| Angles (°) | 2.048 | 2.231 |
| Ramachandran statistics (%) | | |
| Favoured | 95.73 | 96.2 |
| Outliers | 0.32 | 0.48 |

* Statistics for the highest resolution shell are given in parenthesis.

WILEY-VCH

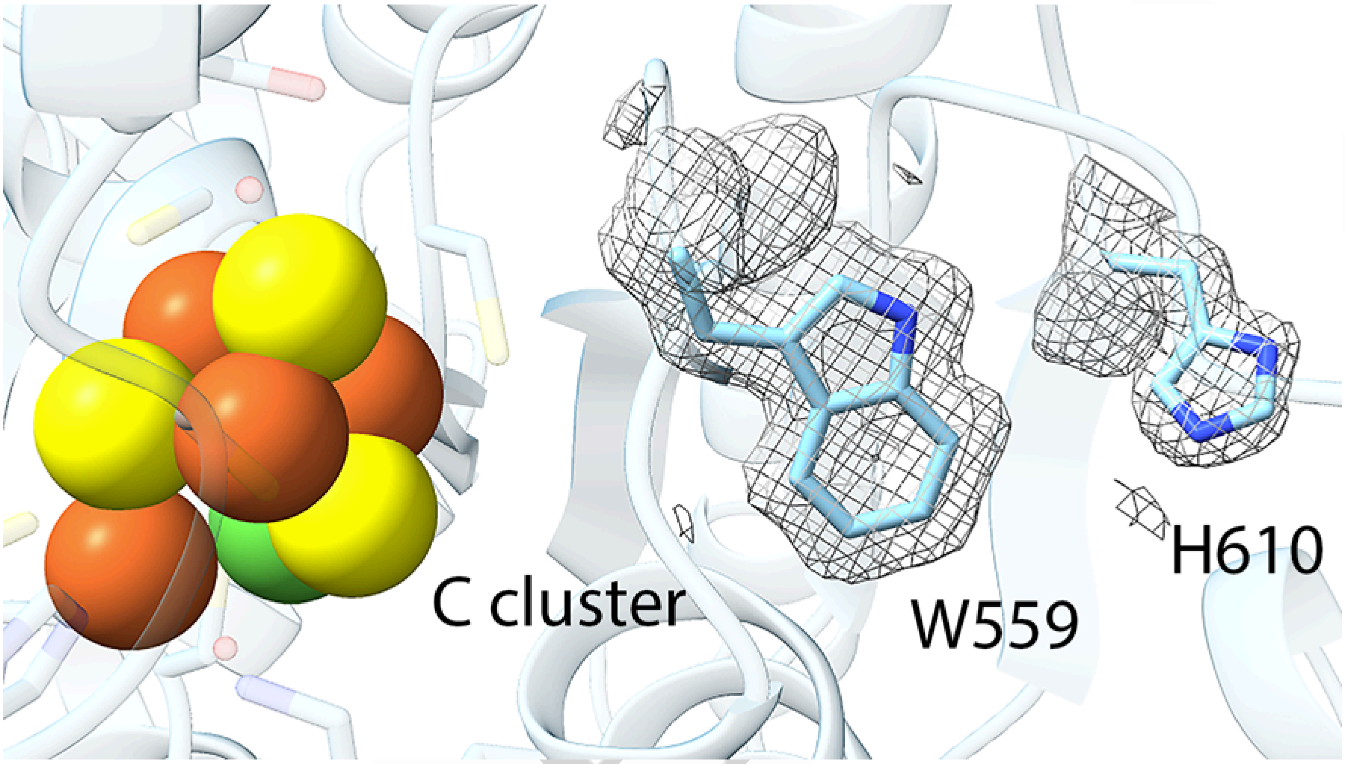


**Supplementary Figure S1.: Omit map of A559W/V610 variant.** The exchanged amino acids A559W and V610H in the double variant of CODH-II$_{Ch}$ are highlighted. The Fo-Fc omit map around the entire residues (sidechain and mainchain) is displayed as grey mesh and contoured at 2 σ.

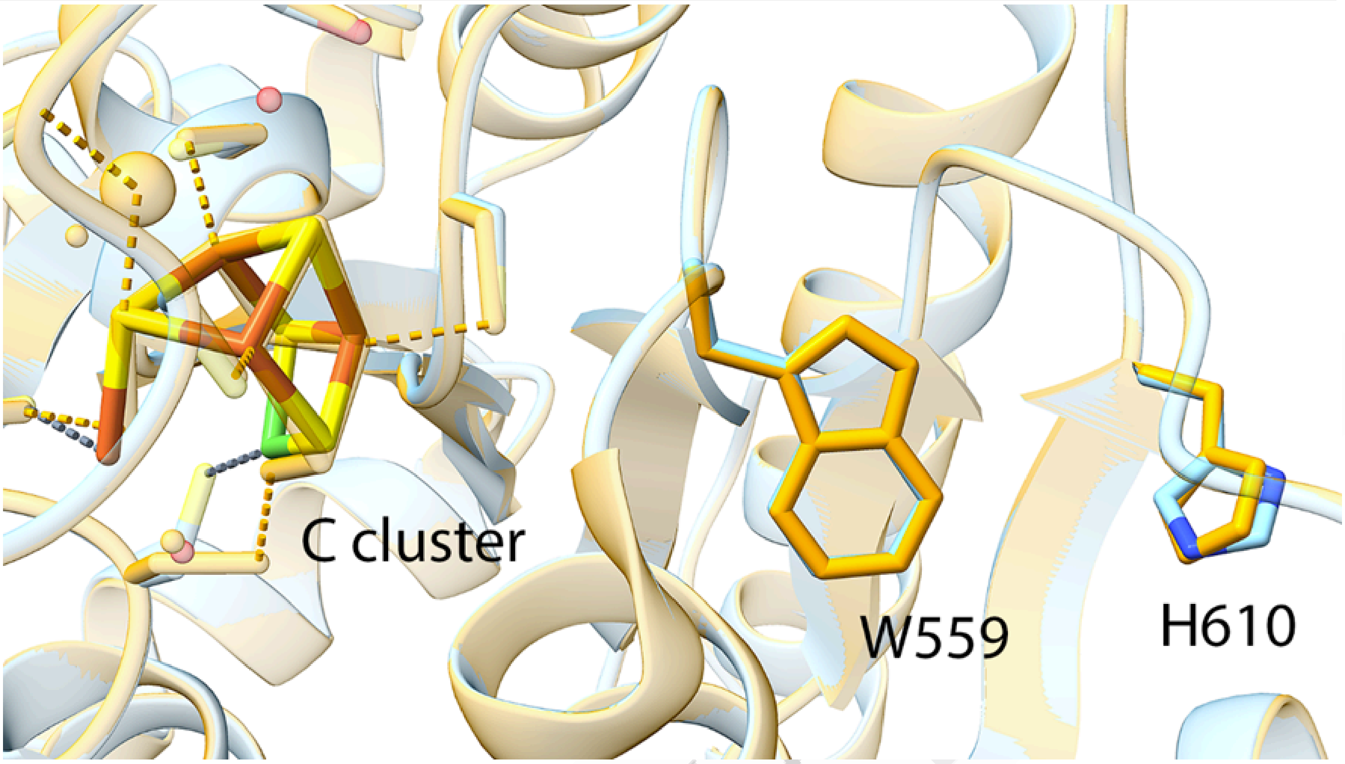


**Supplementary Figure S2.: Comparison of the structure of the double variant A559W V610H reported here with the structure published by Kim et al.** The structure reported of the double variant A559W V610H (depicted in blue) here is superimposed on the structure published by Kim and coworkers[12] (PDB: 9IYM, depicted in orange). The positions of the exchanged amino acids A559W and V610H are highlighted.

### S3 Electrochemistry

We used the approach we previously described to measure the Michaelis constant of the WT and the two mutants, in which the protein film, immobilized on a graphite electrode, is exposed to an injection of CO. A Michaelis-Menten behaviour is fitted to the resulting current, shown in figure S3. Mass transport limitations have to be taken into consideration. The fit yields one value of the $K_M$; an estimation of the error was obtained by repeating the experiments and determining the standard deviation.

We performed our oxygen-experiments using protein film electrochemistry as detailed in [13]. In short, a series of 6 injections of CO was performed in an open electrochemical cell inside a glovebox under constant stirring from a rotating disk electrode. As a consequence, the concentration of injected gas decays exponentially over time. The first two CO injections served to quantify filmloss. The third injection was followed after 10 s by an injection of oxygen, and after the resulting drop in activity we determined the remaining active fraction, which we call “imm”. After oxygen had departed from the solution a 4th CO injection was used to determine the remaining active fraction after oxygen departure, “dep”. A reductive poise was applied going to -560 mV for 20 s, after which a 5th CO injection was made, resulting in the remaining active fraction called “red”. The 6th injection again served to determine film loss.

The electrochemical experiments were performed in an electrochemical cell consisting of a glass jacket permitting water cooling with a platinum wire counter electrode and a saturated calomel reference electrode (Scott instruments) in 0.1 mM NaCl inside a glovebox (Jacomex, France) containing <3 ppm oxygen. The potential was set using a potentiostat from Autolab (PGSTAT128N, Methrohm, the Netherlands) , data was recorded using the GPES software. The potential used in the experiments was -0.31 V vs. SHE. The experiments were performed at 25 °C in a mixed buffer (5 mM MES, 5 mM CHES, 5 mM HEPES, 5 mM TAPS, 5 mM Na acetate, 0.1 M NaCl) at pH 7 under constant stirring of a rotating disk electrode (Princeton applied research, USA) at 4k rpm. The enzyme films were prepared on the rotating disk electrode using simple dropcasting of 0.3

WILEY

µL enzyme from a 0.1-1 mg/ml stock in Tris pH 8 on a pyrolytic graphite edge electrode. CO was injected from a saturated solution in the same mixed buffer prepared by bubbling CO through an 8 mL buffer in a 16 mL Hungate tube closed with a septum for 45 min to remove traces of oxygen. CO was injected using a gas-tight syringe.

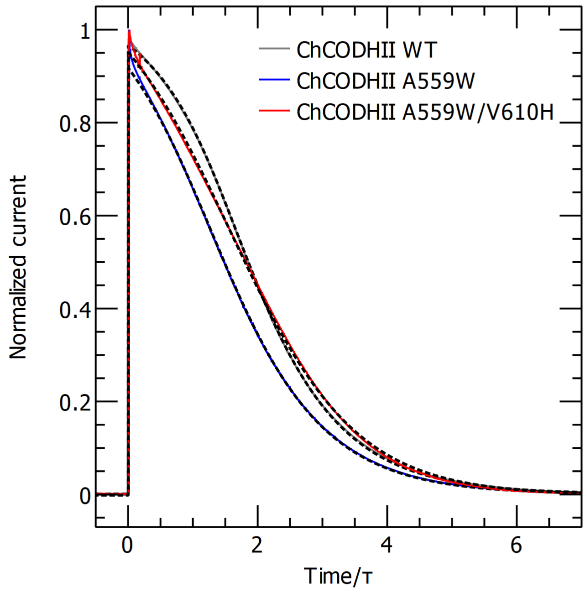


**Supplementary figure S3** experiments to determine the Michaelis constant of *C. hydrogenoformans* CODH II WT, A559W and A559W/V610H

To determine the values of the IC50 and obtain an estimation of the error, we took advantage of the fact that the plot of the remaining activity as a function of injected $O_2$ is almost linear in a log/log plot. Then it is just a matter to fit the equation a(x - x0) to the curve of log10(2 * activity) as a function of log10(O2). The value of x0 is the log10 of the IC50, and the absolute error of the fitted x0 yields the relative error of the IC50 value by using the formula $10^{err}$-1.

WILEY-VCH
